# FUSED: Cross-Domain Integration of Foundation Models for Cancer Drug Response Prediction

**DOI:** 10.1101/2025.09.30.679434

**Authors:** Till Rössner, Jonas Balke, Ming Tang

## Abstract

AI-driven methods for predicting drug responses hold promise for advancing personalized cancer therapy, but cancer heterogeneity and the high cost of data generation pose substantial challenges. Here we explore the transfer learning capability and introduce FUSED (**Fu**sion of Foundation Model **E**mbeddings for **D**rug Response Prediction), a novel architecture for cross-domain foundation model (FM) integration. By systematically benchmark FMs across two domains – molecular FM for drugs and single-cell FM for cell lines, we demonstrate that integrating single-cell FMs substantially reduces the number of input features required for cell line representation. Among FMs, Molformer significantly outperforms ChemBERTa, and scGPT surpasses scFoundation in predictive accuracy and training stability. Moreover, integrating single-cell FMs improves performance in both drug-known and leave-one-drug-out scenarios. These findings highlight the potential of cross-domain FM integration for more efficient and robust drug response prediction.

## 1 Introduction

By leveraging large amounts of pre-training data, Foundation Models (FMs) can represent input in a context beyond the available dataset, making them effective feature extractors for various downstream tasks. In the biomedical domain, pre-trained FMs for medical images [1, 2], drug molecules [3, 4], single-cells [5, 6] and many other modalities are increasingly applied and systematically explored.

Being able to predict drug response is crucial for discovery and design of cancer treatment. The inherently multi-modal nature of the task requires the integration of properties from both drug molecules and cancer cells. Previous published work such as DeepCDR, GraphDRP and XGDP etc. mainly explored various CNN- and GNN-based approaches for extracting and integrating embeddings [7, 8, 9].

In this study, we investigate the potential of FMs for Cancer Drug Response Prediction (CDRP) and present, to our knowledge, the first attempt to integrate FMs across distinct domains. We introduce FUSED (**Fu**sion of Foundation Model **E**mbeddings for **D**rug Response Prediction), and systematically benchmark four representative FMs – scGPT and scFoundation for transcriptomic profiling [5, 6], and ChemBERTa and MolFormer [4, 3] for molecular structure representation. Our work provides a principled path for leveraging cross-domain FMs in robust drug-response prediction.

## 2 Related Work

### Single-Cell Foundation Models

scFMs are pre-trained on large corpora of single-cell gene expression data and have demonstrated capability of representing “virtual cells” and enabling diverse downstream tasks. Among the recent state-of-the-art scFMs, scGPT [5] is a generative pre-trained transformer model built on ~33 million cells, and can effectively distill critical biological insights and achieve superb performance across multiple applications. scFoundation [6], another transformer-based scFM, adopts an asymmetric architecture with a pretraining objective designed to capture complex gene relations across a wide range of cell types and states.

### Molecular Foundation Models

Recent progress in molecular representation learning of chemical drugs has produced FMs that leverage different input modalities [10], including sequence-based models such as ChemBERTa and MolFormer [11, 3], 2D graph-based models such as MolE and KANO [12, 13], and 3D structure-based representations such as Uni-Mol and SubGDiff [14]. In this work, we focus on sequence-based models where drug molecules are represented as SMILES strings. ChemBERTa [11] is pre-trained on 77 million SMILES using masked language modeling to chemical tokens. MolFormer [3] extends this with rotary positional encoding and linear attention.

### Drug Response Prediction

A classical measure of cancer drug response is to estimate the drug efficacy by predicting the half-maximal inhibitory concentration (IC_50_) for drug–cell line pairs. [15]. Most prior work employ hybrid neural network architectures where drug and cell features are encoded separately before their embeddings are integrated. While GraphDRP [8] systematically evaluates different graph neural network (GNN) architectures to encode drug structures, DeepCDR [7] focuses on integrating multi-omics information for cell line representation. More recently, XGDP [9] replaces simple embedding concatenation with a multi-head attention mechanism, yielding more expressive feature interactions than earlier CNN-based fusion.

## 3 Approach

We formulate cancer drug response (CDR) prediction as a regression task, where the model learns a mapping

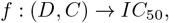

with *D* representing drug features and *C* denoting cell features. Drug features are derived from SMILES representations, while *C* is decomposed into three distinct modalities: transcriptomic, methylation, and mutation profiles. The core idea is to generate embeddings for each of these four input types and integrate them to predict *IC*_50_ values. Our study proceeds in two steps: (1) baseline extension – incorporating FMs into the existing DeepCDR framework to generate drug and cell line embeddings; (2) develop novel integration architecture FUSED (**Fu**sion of Foundation Model **E**mbeddings for **D**rug Response Prediction), which leverages multi-head attention to integrate molecular and single-cell embeddings generated from domain-specific FMs (Fig. 1).

**Figure 1:**
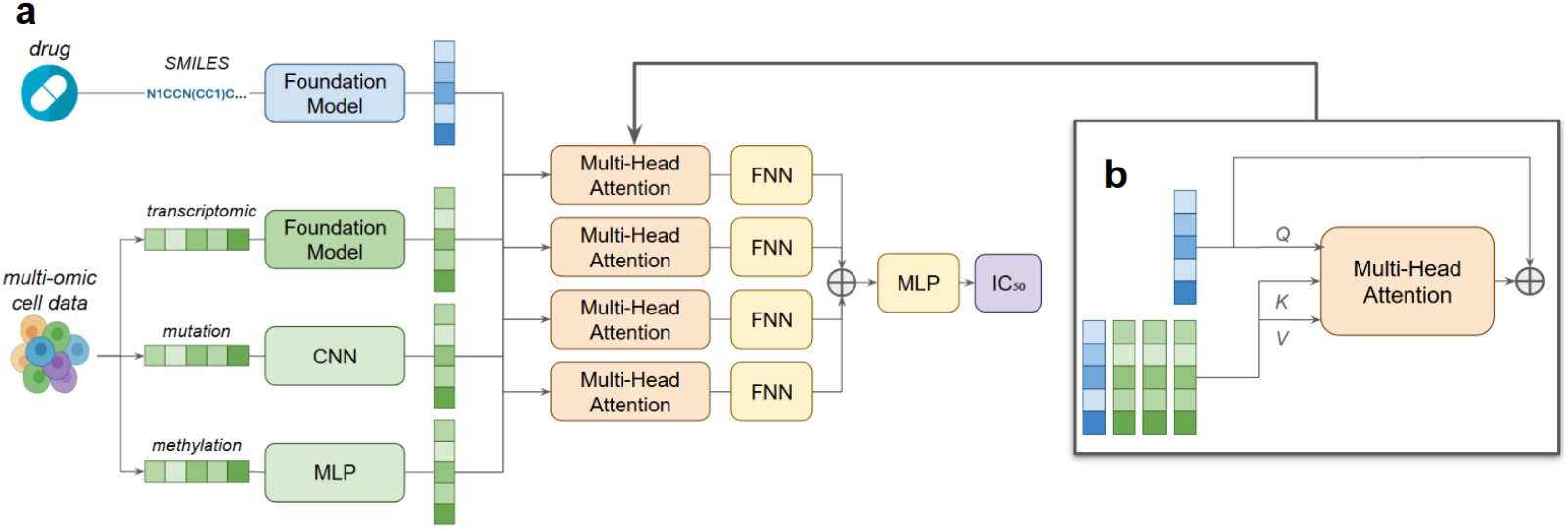
FUSED approach with FMs extracting drug features and transcriptomic features. **a)** We create 4 embeddings for each feature type. While transcriptomics features and drug features are transformed by a FM, mutation and methylation features are trained from scratch. Each embedding is contextualized with respect to the other three embeddings using multi-head attention. All representations are projected to equal dimensions with a linear layer, and concatenated and applied to a MLP to estimate IC_50_. **b)** Multi-head attention mechanism: we create a Q-vector from the original embedding, and a K-, V-vector from all embeddings in order to compute attention score with respect to given Q-vector. Each multi-head attention block learn its own parameters to create Q-, K- and V-vectors.

### Datasets

We use a combination of two common datasets: Cancer Cell Line Encyclopedia (CCLE) [16] and Genomics of Drug Sensitivity in Cancer (GDSC) [17]. CCLE stores multi-omics data (transcriptomics, methylation, mutation) for cancer cell lines. GDSC stores drug responses in form of *IC*_50_ values for cell line drug pairs, matching cell lines from CCLE database. In total, we have 561 cell lines paired with 223 drugs leading to 107446 drug-cell line pairs after filter out missing combinations.

### Architecture Development

Step 1: **Baseline Extension** Our baseline DeepCDR framework contains a graph convolutional network (GCN) and three subnetworks for drug structure and cancer cell profiles (genomic mutation, transcriptomics and methylation) respectively. In our extension, we replace either the GCN-based drug encoder with the molecular FM, or transcriptomic subnetworks with scFM. For the single-cell FM, we prepend it to the transcriptomic-specific subnetwork, while the molecular FM replace the GCN architecture. Step 2: **FUSED** We develop a novel architecture, FUSED, designed to enhance the integration of multiple FMs. Rather than relying solely on concatenation, FUSED employ multi-head attention to contextualize and align embeddings from different modalities before regression (Fig. 1). To optimize hyperparameters (*number of heads, number of MLP layers, learning rate, and embedding dimension*), we performed a grid search in preliminary experiments.

### Experimental Configurations

We used ChemBERTa or Molformer as FMs to extract drug embeddings, and scGPT or scFoundation to extract transcriptomic embeddings. In all settings, FM parameters were kept frozen to assess zero-shot capabilities. All experiments were conducted on an NVIDIA GTX 1080Ti GPU. The training time for the combined multi-omics and GCN model was ~150 minutes (100 epochs). When the GCN component was removed and frozen molecular FMs were used instead, the training time was reduced to ~70 minutes. FUSED required ~10 minutes more time than CNNs.

As primary evaluation metrics, we employed Pearson Correlation Coefficient (PCC, the higher the better) and Root Mean Squared Error (RMSE, the smaller the better). We explored two ways of splitting training/test data. In drug-known split, all drugs and cell lines were present in the training set, and the results are averaged across three runs with different seeds to avoid bias in test set. In leave-one-drug-out split, all samples associated with a given drug were placed exclusively in the test set, ensuring that the model had no prior exposure to that drug during training. This process was repeated for each of the 223 drugs, and the results were averaged across all iterations.

## 4 Results and Discussion

### Drug-Known Split

As shown in Table 1, when all drugs and cell lines appear in the training set, we observe following trends from both PCC and RMSE:

**Table 1:** Performance comparison in drug-known split. Top 3 models from each column are highlighted.

| Drug Encoder + Cell Encoder | Transcriptomics |  | Multi-Omics |  |
| --- | --- | --- | --- | --- |
|  | PCC | RMSE | PCC | RMSE |
| GCN + Subnetworks (DeepCDR) | 0.841 ± 0.003 | 1.483 ± 0.009 | **0.920 ± 0.002** | **1.076 ± 0.007** |
| GCN + scFoundation | 0.882 ± 0.005 | 1.299 ± 0.027 | **0.920 ± 0.003** | 1.076 ± 0.019 |
| GCN + scGPT | **0.914 ± 0.003** | **1.115 ± 0.015** | **0.920 ± 0.000** | 1.076 ± 0.004 |
| ChemBERTa + Subnetworks | 0.821 ± 0.002 | 1.598 ± 0.032 | 0.878 ± 0.003 | 1.326 ± 0.015 |
| ChemBERTa + scFoundation | 0.858 ± 0.006 | 1.440 ± 0.017 | 0.877 ± 0.008 | 1.322 ± 0.027 |
| ChemBERTa + scGPT | 0.876 ± 0.008 | 1.321 ± 0.034 | 0.881 ± 0.006 | 1.332 ± 0.037 |
| MolFormer + Subnetworks | 0.848 ± 0.009 | 1.454 ± 0.037 | **0.919 ± 0.002** | 1.081 ± 0.011 |
| MolFormer + scFoundation | 0.884 ± 0.003 | 1.308 ± 0.004 | **0.919 ± 0.002** | 1.089 ± 0.012 |
| MolFormer + scGPT | **0.912 ± 0.002** | **1.125 ± 0.013** | **0.919 ± 0.003** | 1.084 ± 0.018 |
| FUSED - MolFormer + scGPT | **0.923 ± 0.001** | **1.103 ± 0.094** | **0.932 ± 0.000** | **0.993 ± 0.006** |
| FUSED - MolFormer + scFoundation | 0.899 ± 0.005 | 1.141 ± 0.098 | **0.932 ± 0.001** | **0.992 ± 0.006** |

- **Multi-omics adds modest gains**. Incorporating multi-omics features yields improvements, particularly for models that are weaker with transcriptomics alone. This effect narrows the performance gap among good and poor models, therefore most of our following comparisons will focus on the transcriptomics-only results.
- **scFM helps in transcriptomics only setting**. Both scGPT and scFoundation improve over the baseline, with scGPT consistently outperforming scFoundation in all experiments.
- **Molecular FMs differ**. ChemBERTa underperforms the GCN baseline, whereas MolFormer provides a slight improvement.
- **FUSED is the best**. Our FUSED approach achieves the best performance both in transcriptomic and multi-omics setting.

These findings underscore the value of FMs when comprehensive datasets (e.g., multi-omics) are unavailable. By leveraging knowledge acquired during pre-training, a FM-based approach can compensate for missing modalities. At the same time, FM choice matters: not all models confer benefits (e.g., ChemBERTa), and scGPT is more effective than scFoundation in our setting. Importantly, our FUSED approach demonstrates that integrating molecular and transcriptomic representations through an appropriate fusion mechanism can yield further performance gains, highlighting the promise of cross-domain FM integration for cancer drug response prediction.

### Leave-One-Drug-Out

In the challenging drug-blind setting (Table 2), baseline without scFM performs extremely poorly when only transcriptomic features are used: PCC = 0.076 with GCN and 0.107 with MolFormer as the drug encoder. Interestingly this weakness can be mitigated by adding other omics or using a scFM, which raises PCC into the 0.41-0.45 range and reduces RMSE. Our FUSED approach does not yield additional gains in this setting, indicating the possibility of overfitting to the training drugs.

**Table 2:** Leave-one-drug-out performance comparison of the different approaches.

| Drug Encoder + Cell Encoder | Transcriptomics |  | Multi-Omics |  |
| --- | --- | --- | --- | --- |
|  | PCC | RMSE | PCC | RMSE |
| GCN + Subnetworks (DeepCDR) | 0.076 ± 0.164 | 2.447 ± 1.352 | 0.449 ± 0.137 | 2.409 ± 1.443 |
| GCN + scFoundation | 0.433 ± 0.135 | 2.396 ± 1.440 | 0.443 ± 0.154 | 2.425 ± 1.359 |
| GCN + scGPT | 0.442 ± 0.145 | 2.447 ± 1.352 | 0.439 ± 0.147 | 2.402 ± 1.322 |
| MolFormer + Subnetworks | 0.107 ± 0.168 | 2.400 ± 1.455 | **0.452 ± 0.129** | **2.349 ± 1.406** |
| MolFormer + scFoundation | 0.429 ± 0.137 | **2.323 ± 1.396** | 0.447 ± 0.127 | 2.373 ± 1.395 |
| MolFormer + scGPT | **0.447 ± 0.125** | 2.375 ± 1.396 | 0.448 ± 0.130 | 2.355 ± 1.418 |
| FUSED - MolFormer + scGPT | 0.431 ± 0.148 | 2.444 ± 1.411 | 0.434 ± 0.146 | 2.432 ± 1.358 |

Different than the drug-known setting, PCC and RMSE are not always aligned, and standard deviations are relatively large because the summary statistics are averaged over 223 held-out drugs. This variance indicates that performance depends strongly on which drug is withheld, suggesting that drug-specific properties substantially affect model generalization.

Overall, the weak zero-shot generalization observed in drug-blind setting highlights a key limitation of current AI models for drug response prediction and motivates future work on stronger drug encoders, regularization, and augmentation strategies.

### Training Stability

We examine the validation PCC over training epochs in the drug-known setting (Figure 2). In the transcriptomics-only condition (Figure 2a), scFoundation exhibits pronounced instability—large oscillations and occasional collapses—across runs, whereas scGPT converges quickly and monotonically and the non-FM baseline improves only marginally. Adding additional omics (Figure 2b) markedly stabilizes training: the oscillations diminish and scFoundation’s final performance approaches that of scGPT. These results indicate that scGPT-derived embeddings are robust in low-feature transcriptomic settings, while multi-omics integration mitigates scFoundation’s instability and enables competitive accuracy.

**Figure 2:**
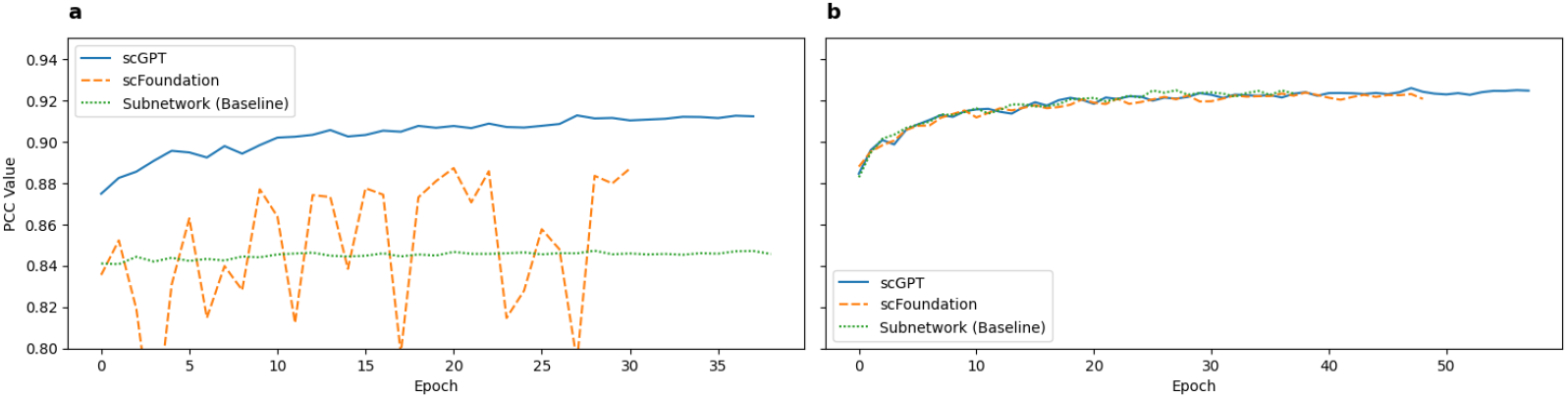
Representative learning curves. PCC validation performance in drug-known setting with MolFormer as drug encoder. a) transcriptomics only; b) multi-omics.

## 5 Conclusion

We introduce FUSED, a compact fusion architecture that aligns molecular and single-cell foundation-model (FM) embeddings, and provide the first systematic cross-domain FM benchmark for cancer drug-response prediction. Across settings, scGPT and MolFormer emerge as the strongest single-modality encoders, and FUSED attains the best accuracy in drug-known splits while reducing reliance on extensive multi-omics features. Limitations include weak generalization to unseen drugs, instability for some scFMs in transcriptomics-only training, and evaluation restricted to frozen encoders on CCLE/GDSC cell-line data. Future work could focus on stronger drug encoders, fusion schemes explicitly regularized for zero-shot drugs, and validation beyond cell lines (e.g. patient-derived models).

## Notes

### Competing Interest Statement

The authors have declared no competing interest.

